# Phenotype bias determines how natural RNA structures occupy the morphospace of all possible shapes

**DOI:** 10.1101/2020.12.03.410605

**Authors:** Kamaludin Dingle, Fatme Ghaddar, Petr Šulc, Ard A. Louis

## Abstract

Morphospaces representations of phenotypic characteristics are often populated unevenly, leaving large parts unoccupied. Such patterns are typically ascribed to contingency, or else to natural selection disfavouring certain parts of the morphospace. The extent to which developmental bias, the tendency of certain phenotypes to preferentially appear as potential variation, also explains these patterns is hotly debated. Here we demonstrate quantitatively that developmental bias is the primary explanation for the occupation of the morphospace of RNA secondary structure (SS) shapes. Upon random mutations, some RNA SS shapes (the frequent ones) are much more likely to appear than others. By using the RNAshapes method to define coarse-grained SS classes, we can directly compare the frequencies that non-coding RNA SS shapes appear in the RNAcentral database to frequencies obtained upon random sampling of sequences. We show that: a) Only the most frequent structures appear in nature; the vast majority of possible structures in the morphospace have not yet been explored. b) Remarkably small numbers of random sequences are needed to produce all the RNA SS shapes found in nature so far. c) Perhaps most surprisingly, the natural frequencies are accurately predicted, over several orders of magnitude in variation, by the likelihood that structures appear upon uniform random sampling of sequences. The ultimate cause of these patterns is not natural selection, but rather strong phenotype bias in the RNA genotype-phenotype map, a type of developmental bias or “findability constraint”, which limits evolutionary dynamics to a hugely reduced subset of structures that are easy to “find”.

## INTRODUCTION

Darwinian evolution proceeds in two steps. First, random changes to genotypes can lead to new heritable phenotypic variation in a population. Next, natural selection ensures that variation with higher fitness is more likely to dominate a population over time. Much of evolutionary theory has focussed on this second step. By contrast, the study of variation has been relatively underdeveloped [1– 13]. If variation is unstructured, or *isotropic*, then this lacuna would be unproblematic. As Stephen J. Gould wrote in a critique of those who make this implicit assumption [3]: *Under these provisos, variation becomes raw material only – an isotropic sphere of potential about the modal form of a species* …*[only] natural selection* …*can manufacture substantial, directional change*.

In other words, if variation is isotropic, then evolutionary trends should primarily be rationalised in terms of natural selection. On the other hand, if there are strong anisotropic developmental biases, then structure in the arrival of variation may well play an important explanatory role in biological phenomena we observe today. While the discussion of how anisotropic variation affects adaptive evolutionary outcomes has moved on significantly from the days of Gould’s critique, primarily due to the growth of the field of evodevo [7], it remains a source of significant contention [5–13].

Unravelling whether a long-term evolutionary trend in the past was primarily caused by biased variation is not straightforward. It often means answering counterfactual questions [14] such as: What kind of variation could have occurred but didn’t due to bias? An important analysis tool for such questions was pioneered by Raup [15], who plotted three key characteristics of coiled snail shell shapes in a diagram called a *morphospace*, finding that only a relatively small fraction of all possible shapes were realised in nature. This concept can be generalised to almost any combination of phenotypic characters [16]. The fundamental reason for the anisotropic occupation of a morphospace could be simply be some form of contingency, where evolution started at one point in the morphospace and did not have enough time to fully explore the space. Or it could be some more predictable cause, such as natural selection disfavouring certain characteristics, or else developmental bias favouring certain types of variation [10, 13].

One way to make progress on these big questions in evolutionary theory is to study genotype-phenotype (GP) maps that are tractable enough to provide access to the full spectrum of possible variation [17, 18], so that counterfactuals [14] can be explored. In this paper, we follow this strategy, employing the well known GP mapping from RNA sequences to secondary structures (SS), to explain in detail how non-coding RNA (ncRNA) found in nature populates the morphospace of all possible RNA SS shapes.

RNA is a versatile molecule. Made of a sequence of 4 different nucleotides (AUCG) it can both encode information as messenger RNA (mRNA), or play myriad functional roles as ncRNA [19]. This ability to take a dual role, both informational and functional, has made it a leading candidate for the origin of life [20]. The number of indentified functional ncRNA types has grown rapidly over the last few decades, driven in part by projects such as ENCODE [21, 22]. Well known examples include transfer RNA (tRNA), catalysts (ribozymes), structural RNA (most famously rRNA in the ribosome), and RNAs that mediate gene regulation such as micro RNAs (miRNA) and riboswitches.

The function of ncRNA is intimately linked to the three-dimensional (3D) structure that linear RNA strands fold into. While much effort has gone into the sequence to 3D structure problem for RNA, it has proven to be stubbornly recalcitrant to efficient solution [23, 24]. By contrast, a simpler challenge, predicting the RNA SS which describes the bonding pattern of a folded RNA, and which is therefore a major determinant of tertiary structure, is much easier to solve [25, 26]. A combination of computational efficiency and accuracy has made RNA SS a popular model for studying basic principles of evolution [25, 27–42]

An important driver of the growing interest in GP maps is that they allow us to open up the black box of variation – to explain, via a stripped down version of the process of development, how changes in genotypes are translated into changes in phenotypes. Unfortunately, it remains much harder to establish how the patterns typically observed in studies of GP maps [17, 18] translate into evolutionary outcomes, because natural selection must then also be taken into account. For GP maps, this means attaching fitness values to phenotypes which is difficult because fitness is hard to measure and is of course dependent on the environment, and so fluctuates.

Progress can still be made by simply ignoring fitness differences, and comparing patterns in nature directly to patterns in the arrival of phenotypic variation generated by uniform random sampling of genotypes, which is also known as ‘genotype sampling’, or *G-sampling* [38]. For example, Smit et al. [31] followed this strategy and found that G-sampling leads to almost identical nucleotide composition distributions for SS motifs such as stems, loops, and bulges as found for naturally occuring structural rRNA. In a similar vein, Jörg et al. [33] calculated the neutral set size (NSS), defined as the number of sequences that fold to a particular structure, using a Monte-Carlo based sampling technique. For the length-range they could study (*L* = 30 to *L* = 50), they found that natural ncRNA from the fR-NAdb database [43] had much larger than average NSS. More recently, Dingle et al. [38] developed a method that makes it possible to calculate the NSS, as well as the distributions of a number of other structural properties, for a much wider range of lengths. They found, for lengths ranging from *L* = 20 up to *L* = 126, that the distribution of NSS sizes of natural ncRNA – calculated by taking the sequences found in the fRNAdb, folding them to find their respective SS, and then working out its NSS using the estimator from [33] – was remarkably similar to the distribution found upon G-sampling. A similar close agreement upon G-sampling was found for several structural elements, such as the distribution of the number of helices, and also for the distribution of the mutational robustness, confirming earlier work on much smaller samples [44].

An alternative to G-sampling is to use uniform random sampling of phenotypes, so called *P-sampling*. If all phenotypes are equally likely to occur under G-sampling, then its outcomes will be similar to P-sampling. If, however, there is a bias towards certain phenotypes under G-sampling, an effect we will call *phenotype bias*, then the two sampling methods will lead to different results. When the authors of [38] calculated the distributions of structural properties such as the number of stems or the mutational robustness under P-sampling, they found large differences compared to natural RNA in the fRNAdb. The fact that G-sampling yields distributions close to those found for natural ncRNA, whereas the counterfactual under P-sampling does not, suggests that bias in the arrival of variation is strongly affecting evolutionary outcomes in nature. As illustrated schematically in Figure 1(a), such a bias towards shapes that appear frequently as potential variation can lead to natural RNA SS taking up only a small fraction of the total morphospace of possible RNA shapes. Here we treat the morphospace more abstractly, but this pattern would carry through with more traditional morphospaces [15] that utilize specific axes to describe phenotypic characteristics or RNA.

**FIG. 1.**
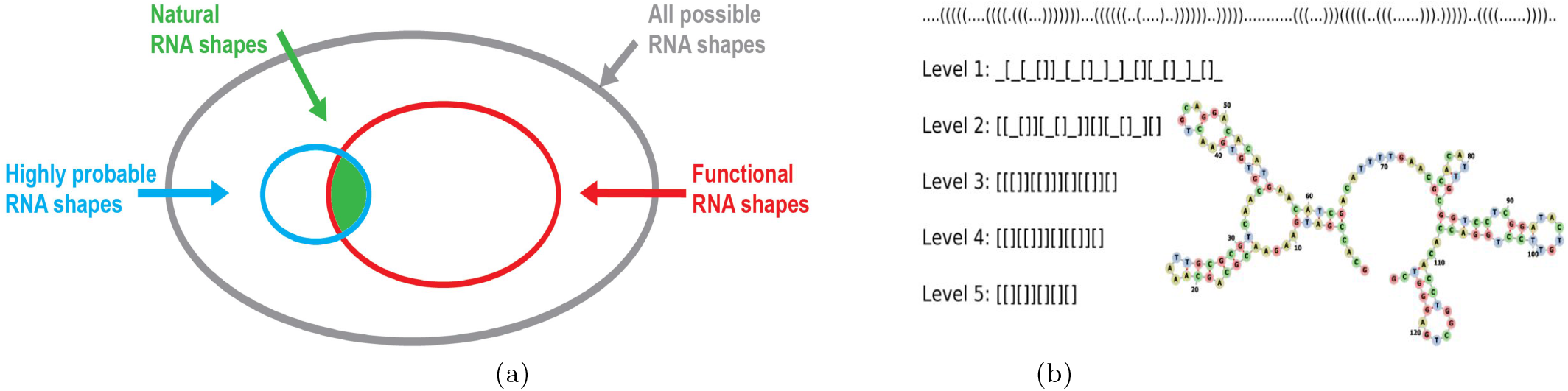
(a) Conceptual diagram of the RNA SS shape morphospace: The set of all potentially functional RNA is a subset of all possible shapes. In this paper we show that natural RNA SS shapes only occupy a minuscule fraction of the morphospace of all possible functional RNA SS shapes because of a strong phenotype bias which means that only highly probable (high frequency) shapes are likely to appear as potential variation. We quantitatively predict the identity and frequencies of the natural RNA shapes by randomly sampling sequences for the RNA SS GP map. **(b) RNA coarse-grained shapes:** An illustration of the dot-bracket representation and 5 levels of more coarse-grained abstracted shapes for the 5.8s rRNA (length *L* = 126), a ncRNA. Level 1 abstraction describes the nesting pattern for all loop types and all unpaired regions; Level 2 corresponds to the nesting pattern for all loop types and unpaired regions in external loop and multiloop; Level 3 is the nesting pattern for all loop types, but no unpaired regions. Level 4 is the helix nesting pattern and unpaired regions in external loop and multiloop, and Level 5 is the helix nesting pattern and no unpaired regions.

Nevertheless, the evidence presented so far for this picture of a strong bias in the arrival of variation has been indirect, and only for distributions over SS structures because individual SS rarely appear more than once in the fRNAdb. Moreover, the measurements have often needed theoretical input, in that they used theoretical estimates for the NSS of individual sequences in the ncRNA databases. To conclusively address big questions related to the role of bias in evolutionary outcomes, a more direct measure is needed.

To achieve this goal of directly measuring frequencies, we first note that any tiny change to the bonding pattern of a full SS, illustrated by the dot-bracket notation in Figure 1(b), means a new SS. In practice, however, many small differences are often found in homologues, suggesting that these differences are not critical to function. To capture this intuition that larger scale ‘shape’ is more important than some of the finer features captured by the full dot-bracket notation, Giegerich et al. [45] defined a 5-level hierarchical abstract representation of SS. At each nested level of description, the SS shape is more coarse-grained, as illustrated in Fig 1(b). By grouping together shapes with similar features, frequencies of ncRNA shapes can be directly measured from a given database. Here, we mainly use the large, popular, and up-to-date RNAcentral[46] database.

In this paper, we show that the the frequency *f*_*p*_ with which abstract shapes are found in the RNAcentral database is accurately predicted by frequencies 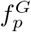 that they are found for G-sampling. We then discuss what these results mean in light of the longstanding controversies about developmental bias.

## RESULTS

### Nature only uses high frequency shapes, which are easily found

We computationally generated random RNA sequences for lengths *L* = 40, 55, 70, 85, 100, 126, and then folded them to their SS using the Vienna package [26], which, as for other closely similar packages based on thermodynamics [47], such as RNAStructure [48] or Unafold [49], is thought to be accurate for the relatively short RNAs we study here (Methods). Next we used the RNA abstract shapes method [45, 50] (See Figure 1(b)), to classify the folded SS into separate abstract structures. Similarly, we also took natural ncRNA sequences from the popular RNAcentral database [46], folded these, and used the RNA abstract shape method to assign structures to them (see Methods).

To compare the G-sampled RNA structures to the natural structures, a balance must be struck between being detailed enough to capture important structural aspects, but not so detailed that for a given dataset very few repeated shapes are found, making it impossible to obtain reliable frequencies. Considering our data sets, we use level 3 for all RNA of length *L* = 40 and *L* = 55 and level 5 for *L ≥* 70. In Figures (S1) and (S2) of the SI we include all 5 other levels for *L* = 55, finding essentially the same results.

Figure (2) shows the shape frequencies 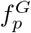 found by G-sampling, ranked from most frequent to least frequent (blue dots). The frequencies, or equivalently the NSS of these structures, vary by many orders of magnitude. The shapes which also appear in the RNAcentral database have been highlighted (yellow circles). Natural ncRNA shapes employ a tiny subset of the most frequent structures. Interestingly, a remarkably small number of random sequences, on the order of 10^3^-10^6^ independent random samples, is enough to find essentially all shapes at these levels of abstraction. found in the RNAcentral database for the lengths studied here. For a sense of scale, there are 4^126^ *≈*7 × 10^75^ sequences of length 126, so that we are sampling on the order of 1 in 10^70^th of the total space and still finding all the structures at the coarse-graining levels chosen. Note that this fraction decreases as the coarse-graining level increases. For example, in Figure (S1) where we show that for *L* = 55 strands, for which there are 4^55^ *≈*10^33^ total possible sequences, we need on the order of 10^7^ sequences for level 1, up to 10^4^ sequences for level 5. These numbers of samples all remain remarkably tiny fractions of the total.

**FIG. 2.**
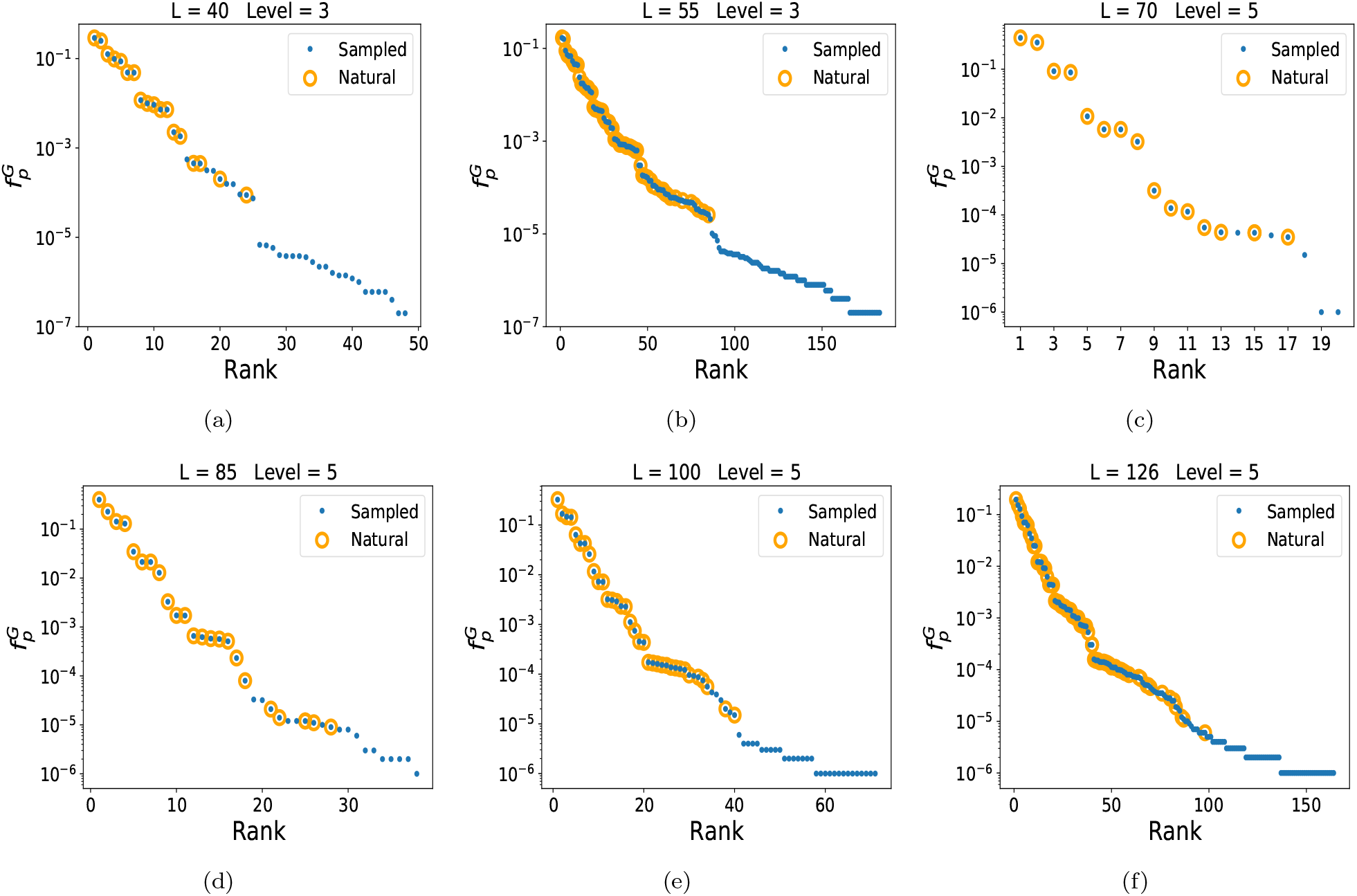
Nature selects highly frequent structures. The frequency 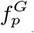 (blue dots) of each abstract shape, calculated by random sampling of sequences (G-sampling), is plotted versus the rank. Yellow circles highlight which of the randomly generated shapes were also found in the RNAcentral database. Panels (a)—(f) are for *L* = 40, 55, 70, 85, 100, 126, respectively. The number of natural shapes are 18, 63, 16, 25, 35, and 68 in order of ascending length, while the numbers of possible shapes in the full morphospace are many orders of magnitude larger, ranging from *≈* 10^4^ possible level 3 shapes for *L* = 40 to*≈* 10^12^ level 5 shapes for *L* = 126. The shapes in nature are all from remarkably small fraction of possible structures that have the highest 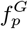 or equivalently the highest NSS. The natural shapes found in the database appear upon relatively modest amounts of random sampling of sequences.

To further quantify just how small a subset of the total morphospace has been explored by nature, we use asymptotic analytic estimates of the total set of possible structures from Table 1 of [51] (but see also earlier results in [52]). These predict 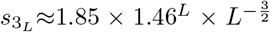 for level 3 and 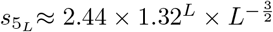 for level 5, where we have taken results pertaining to minimum hairpin length of 3, and minimum ladder length of 1 (which is consistent with the options we used in the Vienna folding package). From these equations we estimate 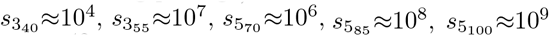, and 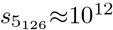. By contrast, in the RNAcentral database we find, at level 3, 18 structures for *L* = 40 and 63 for *L* = 55. At level 5 we find 16, 25, 35, and 68 independent structures for *L* = 70, 85, 100 and 126 respectively. We provide a direct illustration in Figure 3 where the top 183 level 3 structures found for *L* = 55 (after 5 10^6^ sampels) are shown together with the 63 found in nature. The structures employed by natural ncRNA take up an incredibly small fraction of the whole morphospace of possible structures. Moreover, the relative fraction explored decreases rapidly with increasing length.

**FIG. 3.**
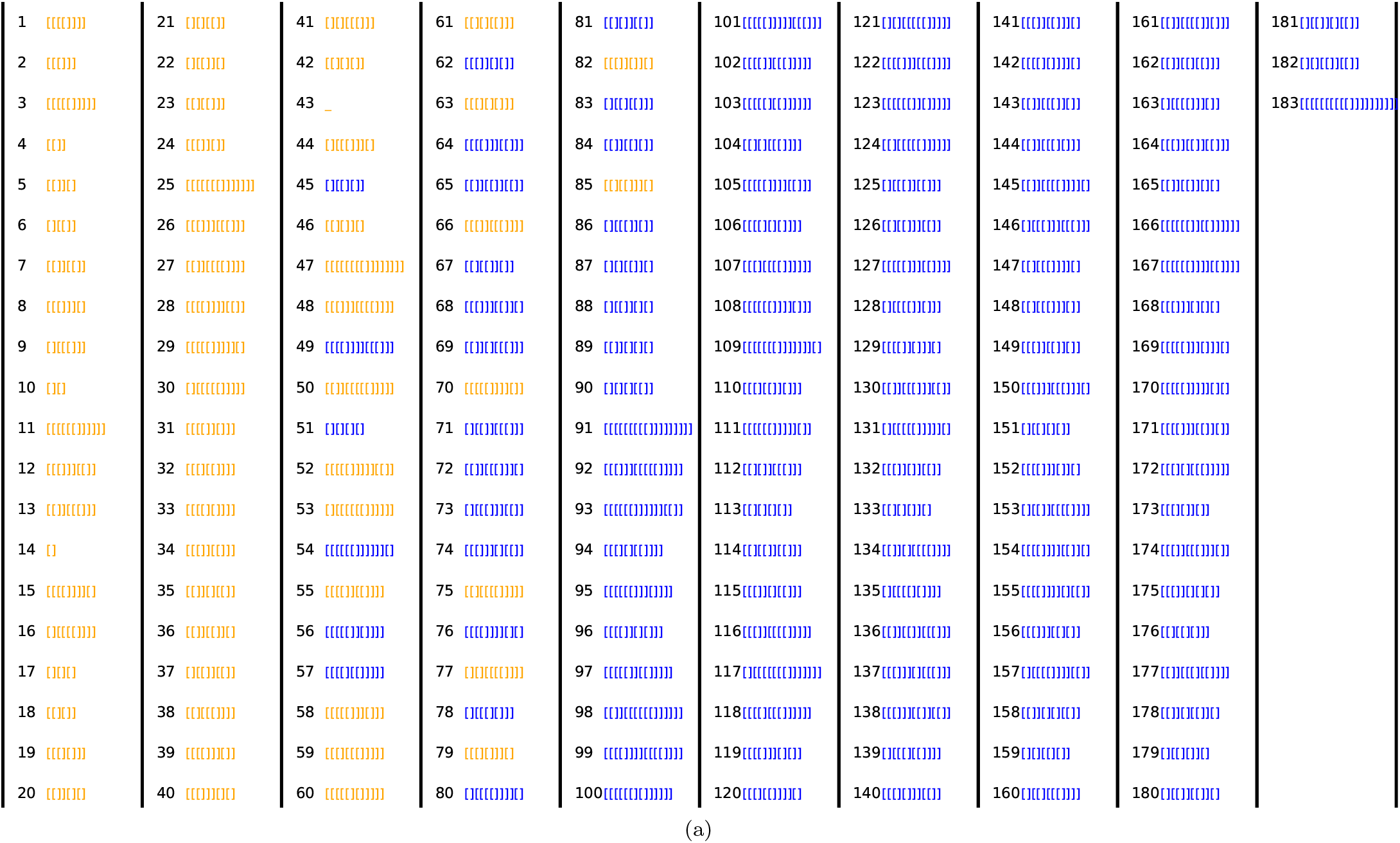
Shape array for L = 55 RNA at level 3, showing the 183 shapes found by sampling 5 × 10^6^ random sequences, in order of their rank by frequency 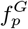. The 63 naturally occurring level 3 shapes from the RNACentral database are highlighted in yellow, demonstrating that only a small fraction of the total morphospace of shapes is occupied by RNAs found in nature, and that these are all highly frequent structures. We estimate that there are on the order of 10^7^ possible level 3 structures for *L* = 55 RNA, so that this array only shows a tiny fraction of the total morphospace of shapes.

### Frequencies of shapes in nature can be predicted from random sampling

Figure (4) demonstrates that the G-sampled frequency of shapes correlates closely with the natural frequency of shapes, for a range of lengths. In SI Figure (S2), we show for *L* = 55 that similar results are found for different levels of shape abstraction, so that this result is not dependent on the level of coarse-graining.

**FIG. 4.**
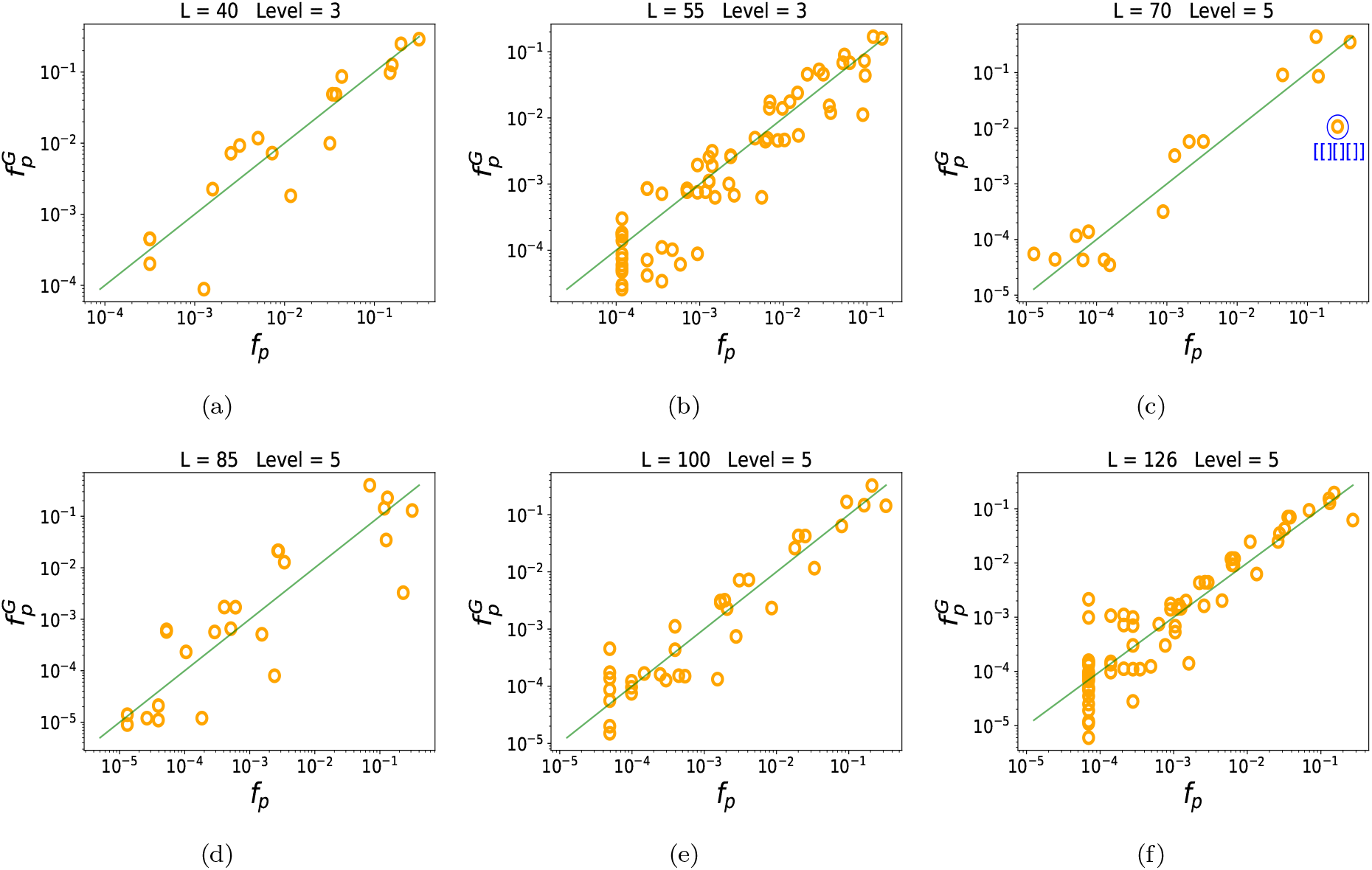
The frequency of shapes in nature correlates with the frequency of shapes from random sampling. Yellow circles denote the frequencies *f*_*p*_ of natural RNA from RNAcentral. The green line denotes *x* = *y*, i.e natural and sampled frequencies coincide. The log frequency upon G-sampling 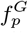 correlates well with *f*_*p*_: (a) *L* = 40 Pearson *r*=0.92; (b) *L* = 55 *r*=0.93; (c) *L* = 70 *r*=0.94; (d) *L* = 85 *r*=0.86; (e) *L* = 100 *r*=0.95; (f) *L* = 126 *r*=0.92; and all correlations have p-value*<*10^*−*6^. We also highlight a blue structure, namely t-RNA for *L* = 70 which has been the subject of extra scientific interest, and is hence over-represented in the database.

We note that there is an important assumption in our interpretation, which is that the frequency with which structures are found in the RNAcentral database is similar to the frequency with which they are found in nature. To first order it is reasonable to assume that this is true, as the databases are typically populated by finding sequences that are conserved in genomes, a process that should not be too highly biased. Moreover, the good correlation between the 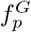 and *f*_*p*_ found here provides additional a posteriori evidence for this assumption as it would be hard to imagine how such close agreement could obtain if there were strong man-made biases in the database. Nevertheless, there are structures that have been the subject of greater researcher interest, and one may expect them to be deposited in the database with higher frequency. We give one example in Figure (4)(c) of an outlier that is over-represented (with high confidence) compared to our prediction, namely the shape [[][][]], which includes the classic clover leaf shape of transfer RNA.

Further, we show in SI section C that qualitatively similar rank and correlation plots (Figure (S3)) appear using the popular Rfam database [53, 54], where structures are determined not by folding, but by a consensus alignment procedure. We also show in SI 5 and SI 6 that our results are robust to changes in CG bias, or when we include other suboptimal structures that are close enough in energy to be accessed by thermal fluctuations. The similar behaviour we find across structure prediction methods, strand lengths, and databases would be extremely odd if artificial biases were strong on average in the natural databases. Hence we believe that our main findings are unlikely to be due to database biases, although at a finer scale there may very well be biases, such as the one we present for tRNA, that are observable, and possibly an interesting source of new insight. Finally, here we have used relatively short RNA for the purposes of computationally tractability and accuracy, in forthcoming work we study much longer RNA, as well as utilizing different methods.

## DISCUSSION

We first recapitulate our main results below under three headings, and discuss their implications for evolutionary theory.

### (A) Nature only utilizes a tiny fraction of the RNA SS phenotypic variation that is potentially available

Besides being an interesting fact about the natural world, this result has implication for synthetic biology as well. There is a vast morphospace [16] of structures that nature has not yet sampled. If these could be artificially created, then they could be mined for new and potentially intriguing functions.

### (B) Remarkably small numbers of sequences are needed to recover the full set of abstract shapes in the RNAcentral database

This effect is enhanced by the fact that we have coarse-grained the SS to allow for direct comparisons. As shown in the SI section A, for finer descriptions of the SS, more sequences are needed to obtain all natural structures, but the numbers remain remarkably small.

For a sense of the scale of the tiny numbers of sequences needed to produce the full spectrum of structures found in nature, consider that the total number of sequences *N*_*G*_ grows exponentially with length as *N*_*G*_ = 4^*L*^. This scaling implies unimaginably vast numbers of possible sequences, even for modest RNA lengths. For example, all individual sequences of length *L* = 77 together would weigh more than the earth, while the mass of all combinations of length *L* = 126 would exceed that of the observable universe [14]. Such hyper-astronomically large numbers have been used to argue against the possibility of evolution producing viable phenotypes, based on the claim that the space is too vast to search through. See the Salisbury-Maynard Smith controversy [55, 56] for an iconic example of this trope. And it is not just evolutionary skeptics who have made such claims. In an influential essay, Francois Jacob wrote [57]:

*The probability that a functional protein would appear de novo by random association of amino acids is practically zero*.

A similar argument could be made for RNA. Our results suggest instead that a surprisingly small number of random sequences is enough to generate all the basic RNA structures needed for life in all its diversity. This finding is relevant for the RNA world hypothesis, since it suggests that relatively small numbers of sequences are needed to facilitate primitive life. In the same vein, it helps explain why random RNAs can already exhibit a remarkable amount of function [58], similarly to what is suggested for proteins in the rapidly developing field of de novo gene birth [59–62].

### (C) The frequency with which structures are found in nature is remarkably well predicted by simple G-sampling

This result is perhaps the most surprising of the three because these G-sampling ignores natural selection. It is widely thought that structure plays an important part in biological function, and so should be under selection.

The key to understanding results (A)–(C) above can be found in one of the most striking properties of the RNA SS GP map, namely strong phenotype bias which manifests in the enormous differences in the G-sampled frequencies (or equivalently the NSS) of the SS [25]. For example, for *L* = 20 RNA, the largest system for which exhaustive enumeration was performed [36], the difference in the 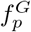 between the most frequent SS phenotype and the least frequent SS phenotype was found to be 10 orders of magnitude. For *L* = 100 this difference was estimated to be over 50 orders of magnitude [38, 41]. Such phenotype bias also explains why G-sampling and P-sampling are so different [38]: a small fraction of high frequency phenotypes take up the majority of the genotypes, and thus dominate under G-sampling.

Evolutionary modelling that takes strong bias in the arrival of variation into account is rare. Population-genetic models that do include new mutations typically consider a genotype-to-fitness map, which often includes an implicit assumption that all phenotypes are equally likely to appear as potential variation, something akin to P-sampling. A notable exception is work by Yampolsky and Stoltzfus [63] which has been applied, for example, to the effect of mutational biases [64, 65].

For the specific case of RNA, however, the effect of strong phenotype bias was treated explicitly in ref [36], where it was shown for evolutionary dynamics simulations ranging from the low mutation monomorphic to the high mutation polymorphic regimes that the mean rate *φ*_*pq*_ at which new variation *p* appears in a population made up of phenotype *q* can be quite accurately approximated as 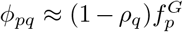, where *ρ*_*q*_ is the mean mutational robustness of genotypes mapping to *q*. In other words, the *average* local rate at which variation *p* appears in an evolving population closely tracks the global frequency 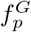, which is exactly what G-sampling measures.

While it is not so controversial that biases could affect outcomes under neutral mutation, see e.g. [66], the strongest disagreements in the field centre around the effect of bias in adaptive mutations [5–13, 64, 65]. Since RNA structure is thought to be adaptive, the main question to answer is how phenotype bias affects RNA evolution when natural selection is also at work. In ref [36], the authors explicitly treat cases where phenotype bias and fitness effects interact. They provide calculations of an effect called the *arrival of the frequent*, where the enormous differences in the rate at which variation arrives implies that frequent phenotypes are likely to fix, even if other higher fitness, but much lower frequency phenotypes are possible in principle. This same effect has also been observed in evolutionary modelling of gene regulatory networks [67]. To avoid confusion, we note that the arrival of the frequent is fundamentally different from the survival of the flattest [68], which is a steady-state effect. There, two phenotypes compete, and at high mutation rates, the one with the largest neutral set size can dominate in a population, even if its fitness is smaller. By contrast, the arrival of the frequent is a non-ergodic effect in the sense that it is not about a steady state with competing phenotypes in a population. Instead, it is about large differences in the rate at which variation appears. Indeed, it can be shown [36] for the RNA GP map, that to first order, the number of generations *T*_*p*_ at which variation on average first appears in a population scales as 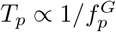 in both the high and the low mutation regimes. Since 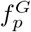 varies over many orders of magnitude, on a typical evolutionary time-scale *T*, only a limited amount of variation (typically that with *T*_*p*_ ≲*T*) can appear. Variation can only fix if it appears in a population. Therefore natural selection acts on SS variation that has been heavily pre-sculpted by the GP map [38].

The close agreement between G-sampling frequencies and measured frequencies of natural ncRNA suggests that once an SS is found that is good enough, natural selection mainly works by further refining parts of the sequence for function, rather than significantly altering the SS. Taken together, these arguments suggest that the arrival of the frequent picture, which is fundamentally about strongly anisotropic variation, provides a mechanism that rationalises all three main classes of observations above.

Nevertheless, given the wide diversity of possible fitness functions that will have played a role in the emergence of the different RNA structures found in the RNACentral database, the argument above that strong phenotype bias determines the outcomes of evolutionary dynamics in such a predictable way may still seem quite surprising. There is, however, a fascinating connection between the arrival of the frequent effect in evolution, and a related behaviour in the dynamics of optimisation in deep neural networks (DNNs). One way of thinking about DNNs is as mappings from the (adjustable) parameters of the DNN (the genotypes), to functions (the phenotypes), which describe how input data maps to a DNN output. The volume of parameters mapping to a particular function is directly proportional to the probability that this function obtains upon random sampling of parameters, and is analagous to the NSS in GP maps. As was found for the RNA GP maps, the mapping from parameters to DNN functions can be hugely biased [69, 70].

Of course DNNs are not trained by randomly sampling parameters, just as an evolutionary process does not use G-sampling either. Instead, the most popular way to optimise DNNs is by using stochastic gradient descent (SGD) [71] which follows the contours of a complex loss-landscape, much as evolution follows a fitness-landscape over time. For such highly biased systems, the arrival of the frequent phenomenology predicts that functions with a large volume of parameters mapping to them are much more likely to be found by an optimiser than functions with a smaller volume of parameters are. Interestingly, it was recently shown for several DNNs and datasets [72] that the probability that the SGD optimiser converges on a particular function is well approximated by the (Bayesian) probability that this function obtains upon random sampling of DNN parameters, which is directly analogous to G-sampling. Since this phenomenology was observed for multiple systems and loss functions, it suggests that that a mechanism much like the arrival of the frequent works robustly for these highly biased systems also. As for the RNA system, where G-sampling provides a good first-order prediction of the frequencies of RNA structures found in nature, so for DNNs, random sampling of parameters provides a good first-order prediction of the frequencies that a DNN converges to a particular function when it is optimised by SGD. This analogy between optimisation on a loss landscape for DNNs and evolutionary dynamics on fitness landscapes strengthens our hypothesis that the arrival of the frequency mechanism can explain why, for highly biased GP maps, multiple evolutionary scenarios produce outputs with probabilities given by G-sampling.

It is interesting to consider whether our arguments that strong phenotype bias affects adaptive evolution can shed light on a related controversy around mutational biases. For example Stoltzfus and McCandlish [64], argued that transition-transversion mutation bias in the arrival of mutations can affect the frequency of adaptive amino acid substitutions. This conclusion was criticised by Svensson and Berger [11], who argued that the bias may not be large enough to overcome fitness differences, and that there may be alternative adaptive arguments for the codon substitution patterns observed in [64]. The basic arguments behind mutational biases having an effect in adaptive evolution are similar in spirit to our arguments for phenotype bias, but there are also differences. Phenotype bias is about the rate at which phenotypes arise, and here we treat all mutations as being equally likely, while mutational bias captures inhomogeneities in the rate at which mutations arise along a genome.

Mutation bias is also typically much smaller than phenotype bias [64, 65, 73]. The global differences in the 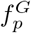 are enormous. Even in the presence of adaptive forces, this “findability constraint” limits the evolutionary process to a tiny subset of high frequency phenotypes in the morphospace. Within the subset of phenotypes that are found, however, the relative differences in frequencies are relatively small (on the order of 4 to 5 orders of magnitude in range). As long as phenotypes are findable, one might think that adaptive forces could overwhelm the developmental bias, which manifests in differential rates in the arrival of variation [5, 8, 11]. Interestingly, we nevertheless observe a fairly close correlation between *f*_*p*_ and 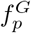 which suggests that, averaged across many evolutionary scenarios, relatively small differences in the arrival of variation, such as those expected under mutational bias [64, 65, 73], may indeed affect adaptive evolution.

Strong phenotype bias is also consistent with SELEX experiments [74, 75], where artificial selection for RNA function can, with a relatively small amount of material, lead to the repeated convergent evolution of the same structures. Famous examples include RNA aptamers [76] and the hammerhead ribozyme [77], which also shows convergence in nature [62]. In light of the unimaginably small portion of the hyper-astronomically large sequence spaces these experiments explore, this convergent evolution seems highly surprising. But when we consider the strong phenotype bias, then a possible explanation emerges. SELEX experiments rely on artificial selection to refine sequences and hone in on a particular function. While natural selection is the ultimate reason why a particular *function* emerges (such as self-cleaving catalytic activity for the hammerhead ribozyme), we hypothesise that the same *structures* emerge because of phenotype bias. After all, multiple structures could, in principle, produce the same function. In other words, to use Mayr’s famous ultimate-proximate distinction [78–80] for RNA SS, phenotype bias is the ultimate, and not merely the proximate cause of the evolutionary convergence of the structures found in SELEX experiments and in nature. The idea that developmental biases could help explain convergence is not new, but we believe that the type of phenotype bias we are proposing here has not yet been seriously considered as a cause of convergence.

How is phenotype bias related to the broader literature on developmental bias? To first order, phenotype bias is just another way of expressing developmental bias: certain phenotypes are more likely than others to appear upon mutations. Early work mainly considered developmental bias as constraint [1], in that it limits what kind of variation natural selection can work on. The phenotype bias we observe can be viewed that way. Whether it can also act as a developmental drive [81] that facilitates adaptive evolution would hinge on there being advantages to the kinds of structures that it favours. Indeed, G-sampled RNA structures are on average different from P-sampled structures. For example they have higher mutational robustness, and fewer stems [38]. So there is bias towards these characteristics, which may be adaptive.

Where RNA phenotype bias differs the most from classic examples of developmental bias such as the universal pentadactyl nature of tetrapod limbs, is that the latter are thought to occur because evolution took a particular turn in the past that locked in a developmental pathway, most likely through shared ancestral regulatory processes [13]. If one were to rerun the tape of life again, then it is conceivable that a different number of digits would be the norm. By contrast phenotype bias predicts that the same spectrum of RNA shapes would appear, populating the morphospace in the same way. It is true that given enough time, a larger set of RNA shapes could appear, but the exponential nature of phenotype bias implies that orders of magnitude more time are needed to see linear increases in the number of potential shapes, so that we can be pretty confident that a broadly similar spectrum of shapes would appear again.

It is also interesting to compare phenotype bias to adaptive constraints. For example, there are many scaling laws observed in nature. One of the most dramatic is Kleibers law which states that the metabolic rate of organisms scales as their mass to the 3*/*4 power, and which has been shown to hold over a remarkable 27 orders of magnitude [82]! The morphospace of metabolic rates and masses is therefore highly constrained. Such scaling laws can be understood in an adaptive framework from the interaction between various basic physical constraints [82], rather than from biases in the arrival of variation. Phenotype bias also arises from a fundamental physical process [83] and limits the occupation of the RNA morphospace. But it provides, by contrast, a non-adaptive explanation for the constraint. At this level, it may be closest in spirit to some constraints that are postulated in biological or process structuralism [84] since the phenotype bias “findability constraint” arises from the GP map itself.

In the literature, nomenclature around developmental biases and evolutionary constraints is not completely settled, and that ambiguity affects our discussions above. Phenotype bias is interesting in this regard because on a larger scale it is perhaps most naturally described as a findability constraint, while on the smaller scale of those phenotypes that are found, the close agreement between *f*_*p*_ and 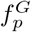 is perhaps most naturally described as a developmental bias or a developmental drive.

Finally, the fact that G-sampling does such a good job at predicting the likelihood that SS structures are found in nature also has implications for the study of selective processes in RNA structure [85, 86]. We propose here that signatures of natural selection should be measured by considering deviations from the null-model provided by G-sampling. Our current work has been on relatively short sequences, where simple SS folding algorithms based on thermodynamics are thought to work reasonably well. For longer sequences, other more sophisticated methods that include, for example, information from evolutionary covariance [87], may be be needed.

In conclusion, while the RNA sequence to SS map describes a pared down case of development, this simplicity is also a strength. Just as in the fields of chemistry and physics, where the hydrogen atom provides an important model system because it is so easily solvable, so the RNA SS GP map could be viewed as the “hydrogen atom of developmental biology”. The fact that it is so tractable allows us to explore counterfactual questions [14] such as: what kind of phenotypic variation is possible in principle, but did not appear due to phenotypic bias. This system thus provides, to our knowledge, the cleanest evidence yet for developmental bias strongly affecting evolutionary outcomes.

Many other GP maps also show strong phenotype bias [18, 83]. An important question for future work will be whether there is a universal structure to this phenotype bias that holds more widely and whether it also has such a clear effect on evolutionary outcomes in other biological systems. In this context, we note a recent proposal [88] that applies a result related to the coding theorem of algorithmic information theory (AIT) [83] to predict that GP maps should be generically biased towards phenotypes with low descriptional (Kolmogorov) complexity. In close analogy to what we found for natural RNA here, such simplicity bias was demonstrated directly for protein quaternary structures in nature as well as for a related polyomino model for protein quaternary structures [89–91], and also in a gene regulatory network. If it is indeed the case that strong simplicity bias is common in nature, and if, as also suggested in [88], the arrival of the frequent mechanism is important for the evolutionary dynamics of this much wider set of systems, then the conclusions for evolutionary causation driven by strong phenotype bias we draw here for RNA should hold much more widely in nature.

## MATERIALS AND METHODS

### Folding RNA

We use the popular Vienna package [26], based on the Turner model thermodynamics [47], to fold sequences to structures, with all parameters set to their default values (e.g. the temperature *T* = 37^*°*^*C*). This method, much like others in its class, is thought to be especially accurate for shorter RNA. The numbers of random samples were 5×10^6^ for *L* = 40 and *L* = 55, and 10^6^ for *L* = 70, 85, 100, 126. For G-sampling, we choose random sequences, and fold each one. Sequences from the RNAcentral database were folded using the Vienna package with the same parameters as above, after removing any duplicate sequences.

### Abstract shapes

RNA SS can be abstracted in standard dot-bracket notation, where brackets denote bonds, and dots denote unbonded pairs. To obtain coarse-grained abstract shapes [50] of differing levels we used the RNAshapes tool available at https://bibiserv.cebitec.uni-bielefeld.de/rnashapes and the Bioconda rnashapes package available at https://anaconda.org/bioconda/rnashapes. The option to allow single bonded pairs was selected, to accommodate the Vienna folded structures which can contain these.

### Natural sequences

For each length, we took all available natural non-coding RNA sequences from the RNAcentral database [46]. Any repeated sequences were discarded. The sequence secondary structure predictions were made by the Vienna package, and then abstract RNA shapes for each structure were obtained.

The numbers of natural sequences, numbers of shapes, and fractions of natural shapes found by random sampling, were:

*L* = 40: 3160 sequences, yielding 18 unique shapes at level 3 (18/18 shapes were found by random sampling);

*L* = 55: 8619 sequences, yielding 63 unique shapes at level 3 (63/63 found);

*L* = 70: 78075 sequences, yielding 16 unique shapes at level 5 (16/16 found);

*L* = 85: 76143 sequences, yielding 25 unique shapes at level 5 (24/25 found);

*L* = 100: 20314 sequences, yielding 35 unique shapes at level 5 (35/35 found);

*L* = 126: 14318 sequences, yielding 68 unique shapes at level 5 (68/68 found).

In total 224/225 (ie *>*99%) of the shapes in the database were found by relatively modest sampling.

## ACKNOWLEDGMENTS

We thank David McCandlish for discussions and for suggesting the H-atom analogy for RNA GP maps as models for development, and Michael Matthies for help with Fig 1(a).

## Supplementary Information for

### A. *L* = 55 data for levels 1 to 5

In Figures (S1) and (S2) we show plots for the *L* = 55 data using all five coarse-grained abstraction levels of RNAshapes from Giegerich et al. [1]. These figures demonstrate very similar results to those found in the main text for level 3. This qualitative agreement strongly suggests that our main findings are robust to our choice of level. Note that the lowest possible frequencies directly measured in the database are limited by the relatively small number of samples, which affects lower levels of coarse-graining more strongly, because there are more such shapes available. The rank plots in Figure (S1) suggest that as more sequences are added, a wider range of frequencies will be found, improving the correlation at low frequency in Figure (S2).

### B. *L* ≈ 100 data from Rfam

To briefly check that our results maintain for a different database, and with secondary structures not obtained solely by folding algorithms, here we study data from the Rfam [2, 3] database.

All RNAs of length 95 to 105 were taken from all available seed sequences of ncRNA families from the Rfam database. Their secondary structures were obtained by aligning to the consensus structure of the seed alignment for respective RNA families. Note that this is different to analysis we performed for the main text, where instead secondary structures were predicted via folding algorithms, using the popular Vienna package.

The total number of sequences obtained were 4309, but a small fraction (ie 185 or 4.3%) of these were discarded because they were invalid secondary structures according to the folding rules used by the shape abstracter. For example, some of the consensus structures contained motifs with a loop of length 1 — ie (.) — which are deemed invalid. The reason we combined data for lengths 95 to 105 (rather than just using *L* = 100) is that there were relatively few sequences and RNA shapes for just *L* = 100, and so by combining data from other lengths close to 100, we obtain better statistics.

Qualitatively similar rank and correlation plots appear when using Rfam data for *L* ≈ 100 in Figure (S3) as compared to the correlation plots in the main text. These results provide evidence that the correlations we find between random sampling and the natural RNA are not artefacts of either the database which we have used, nor of the method for obtaining secondary structures.

### C. Effects of GC content

We also examine the effects of altering the GC content of the sequences on the probabilities of RNA shapes. In the natural sequences which we have used, there does not appear to be strong deviations from 50% GC content: the average GC content is lowest at 44% for *L* = 55 and highest at 57% for *L* = 85. Nonetheless, because GC content can affect RNA structures and shapes[4], we here perform some further analysis regarding the role of GC content in our current study.

We performed 1 million samples of *L* = 55 sequences using level 3 abstraction with 50% GC content, ie using *P*(*G*) = *P*(*C*) = *P*(*A*) = *P*(*T*) = 0.25 at each nucleotide base, which yielded 150 different RNA shapes. A separate 1 million random sample at 70% GC content (ie *P*(*G*) = *P*(*C*) = 0.35 and *P*(*A*) = *P*(*T*) = 0.15), yielded 189 shapes. Finally, another separate random sample with 30% GC content (ie *P*(*G*) = *P*(*C*) = 0.15 and *P*(*A*) = *P*(*T*) = 0.35) yielded only 82 shapes.

In Figure (S4) we provide correlation plots showing the probability of abstract shapes under different GC content values, namely 50% GC content vs 30% in (a) and 50% GC content vs 70% in (b). The linear correlations were very high: *r* = 0.95 for 30% (p-val*<* 10^*−*6^) and *r* = 0.97 for 70% (p-val*<* 10^*−*6^). The main effect of the CG content is that neutral set sizes (NSS) are larger for lower CG content, a result that was also obtained in the supplementary materials of [5]. This effect explains why under a fixed number of samples, more structures are observed for higher CG content. However, the relative frequencies are quite similar.

**FIG. S1.**
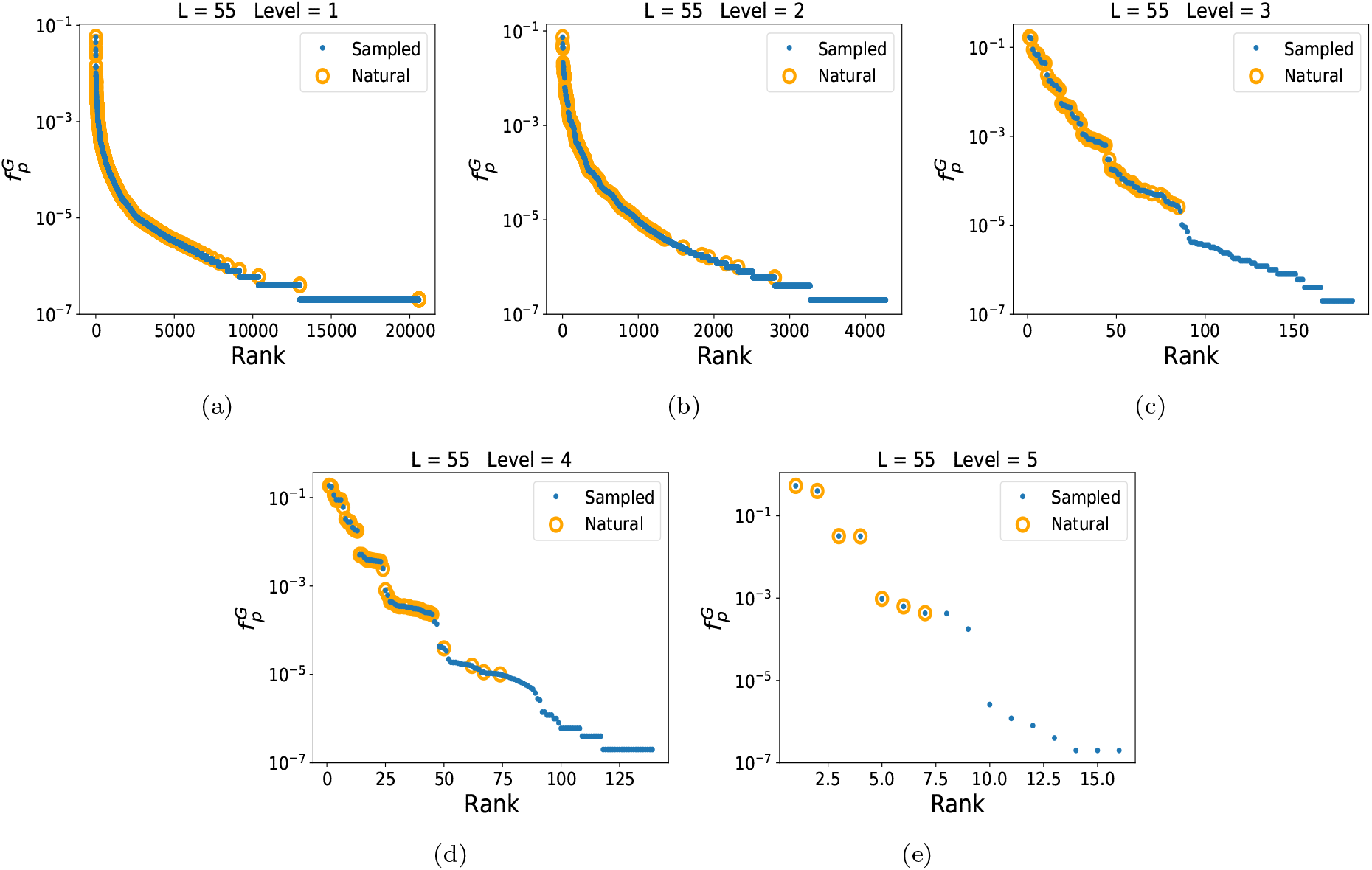
Rank plot for *L* = 55, across all abstraction levels 1, 2, 3, 4 and 5, with 5×10^6^ random samples for each level, compared to the natural frequencies from the RNAcentral database. The number of random shapes and number of natural shapes (in brackets) found for levels 1—5 are 20587 (1083), 4268 (394), 183 (63), 139 (46), and 16 (7).

**FIG. S2.**
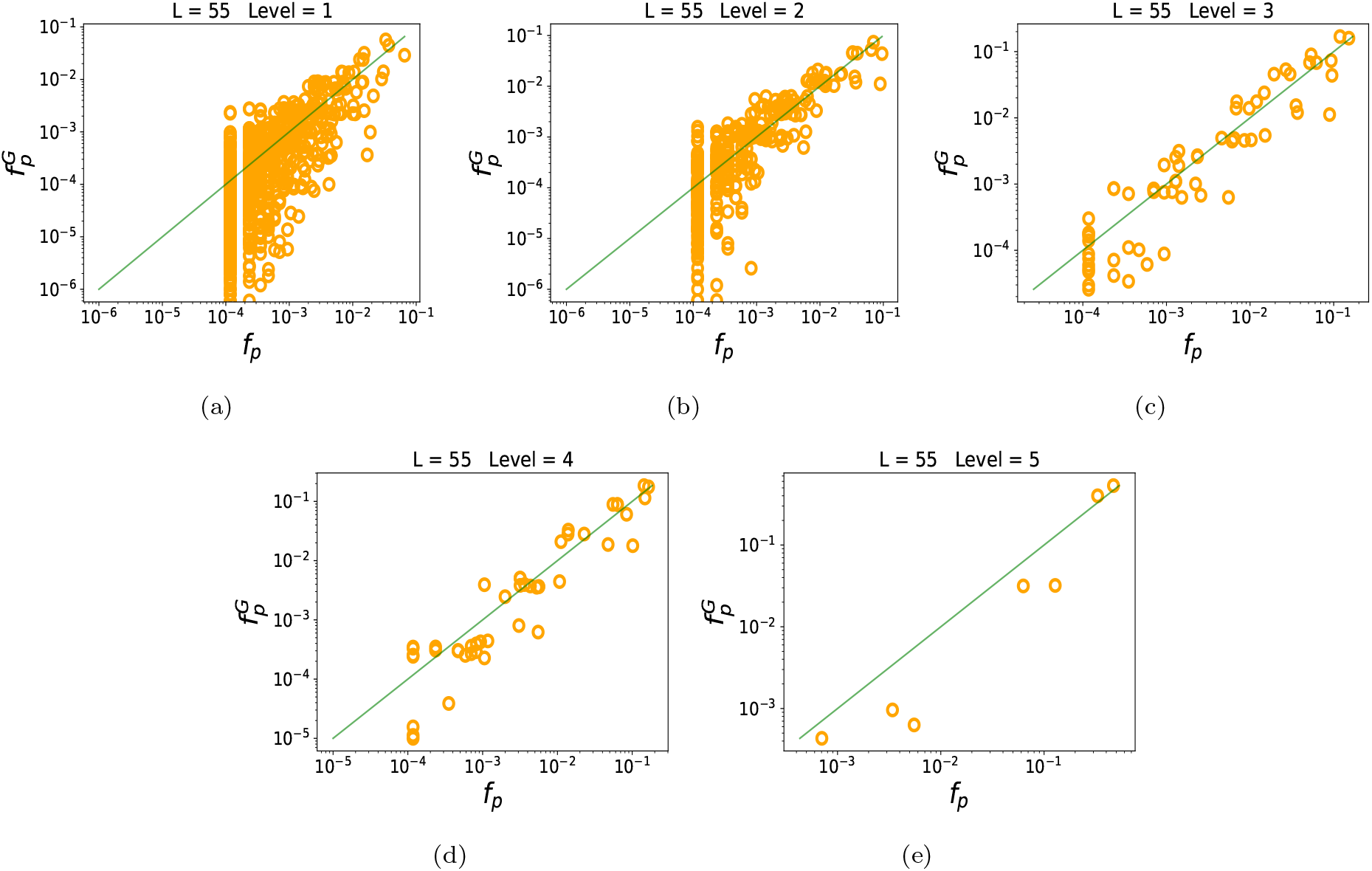
The frequency of shapes in a database correlates with the frequency in nature for *L* = 55, across all abstraction levels 1, 2, 3, 4 and 5, with 5×10^6^ random samples for each level. For lower abstraction levels, there are fewer samples per shape, and hence more noise. With higher levels and hence more samples per shape, there are less points, but also less noise and a clearer correlation. The green line is simply *x* = *y*; it is not a fit to the data.

**FIG. S3.**
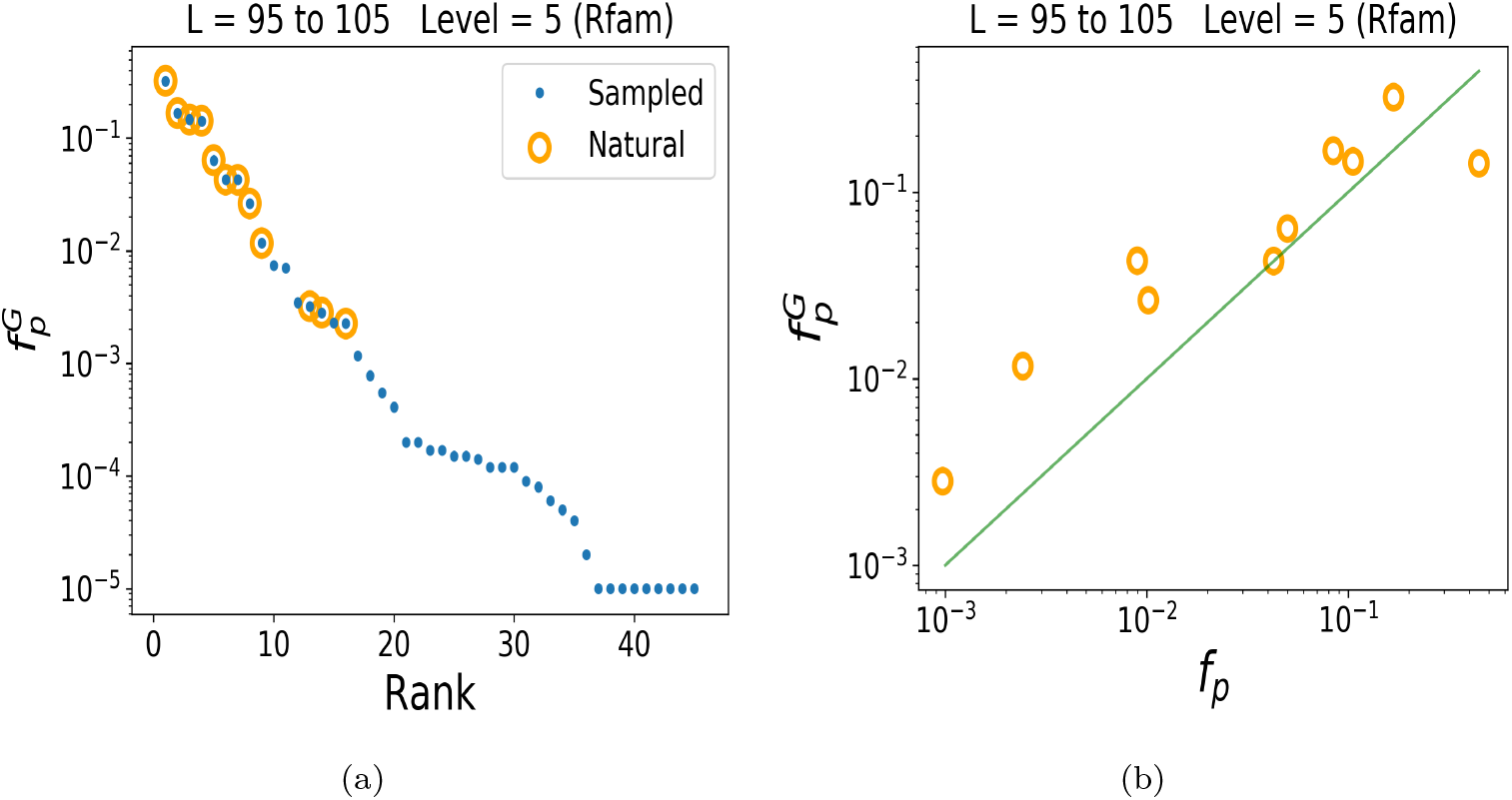
Rank and correlation plots for natural and random data, using Rfam data. (a) Combined data for *L* = 95, 96, …, 104, 105 natural consensus structures rank plot; and (b) *L* = 95 to 105, correlation plot with *r* = 0.96, *p*-value *≈* 10^*−*6^. The data contains 4124 sequences, which yielded 13 unique shapes (level 5).

**FIG. S4.**
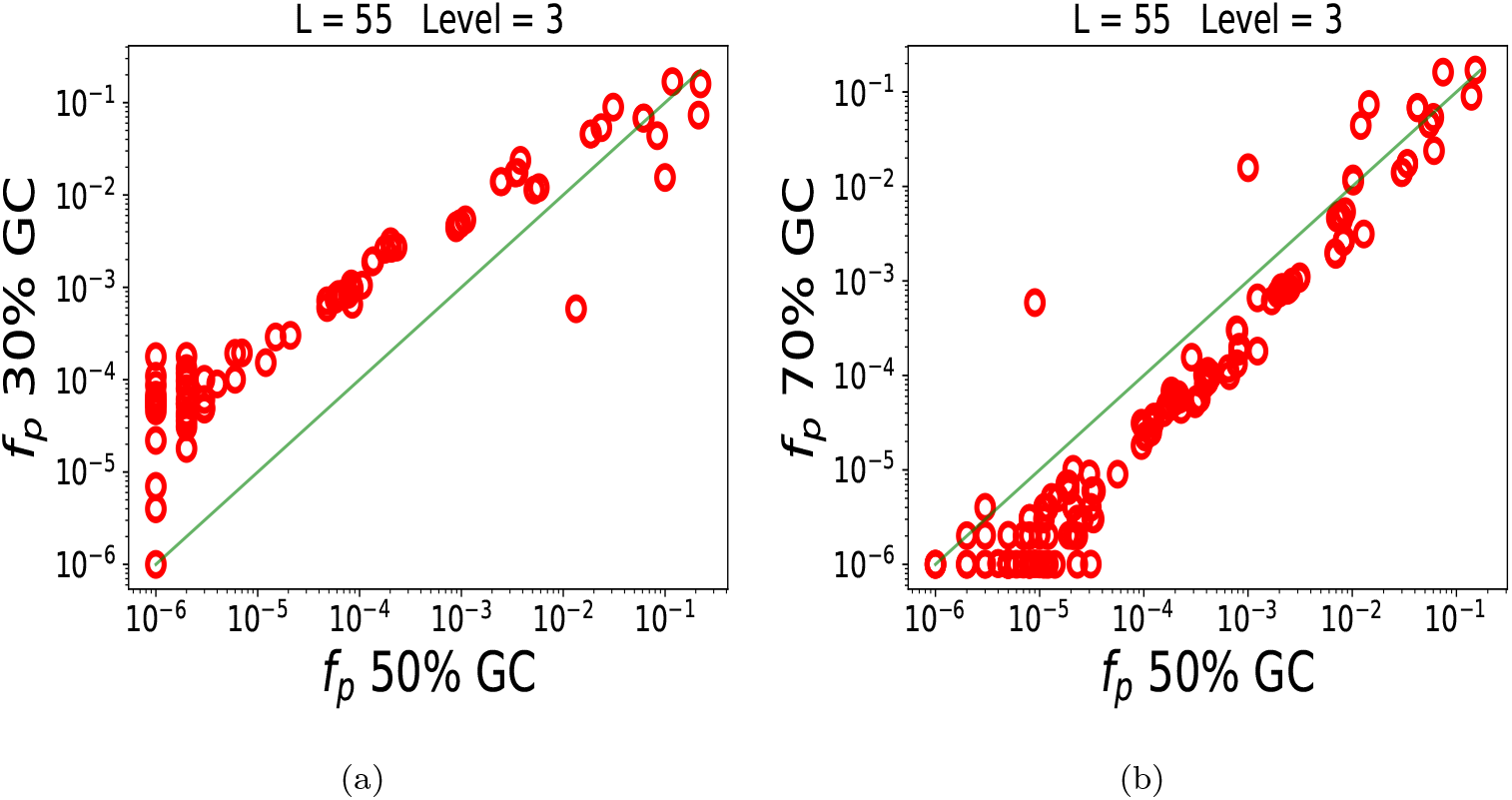
There is a high correlation between the frequencies of RNA shapes when sampling random sequences of different GC contents. (a) 50% GC content vs 30% GC content and (b) 50% vs 70%.

### D. Suboptimal structures

Biologists usually operate with the approximation that there is only one secondary structure for any given RNA sequence. However, if one uses the thermodynamic cost function of a typical folding program, then there are many different RNA structures (hence abstract shapes) which a given sequence can adopt which are within a few *k*_*B*_*T* of the minimum free energy (MFE) structure. Thus one can calculate the fraction of time spent in each shape determined by the free energy of each structure via the Boltzmann distribution. Here we investigate these so-called suboptimal structures, by comparing the probability of abstract shapes from sampling, when a single shape is assigned to a sequence, and the probability of a shape if suboptimal shapes are also included. Suboptimal shapes were calculated using the RNAsubopt function of the Vienna package, see www.tbi.univie.ac.at/RNA/RNAsubopt.1.html.

We study here the correlation between two variables: *f*_*p*_ is the probability of obtaining a given abstract shape on random sampling of a sequence, assuming that only one shape is assigned to each sequence which corresponds to the MFE dot-bracket secondary structure. For example, if a sequence has (((…)))…………. as its MFE structure, then the corresponding shape (level 3) would be [].

The second variable *f*_*p*_ *subopt*., is the probability of obtaining a shape, summing over the Boltzmann probability weights from different sequences. For example if a sequence has (((…)))…………. as its MFE structure, then the corresponding shape (level 3) would be [], but at the same time that sequence would have a certain probability of adopting many different dot bracket structures like eg (((…)))…….(((…))) which would have shape [][]. Summing the Boltzmann weighted probabilities of the shape [][] from different sequences gives the total *f*_*p*_ *subopt*. value for [][] etc…

**FIG. S5.**
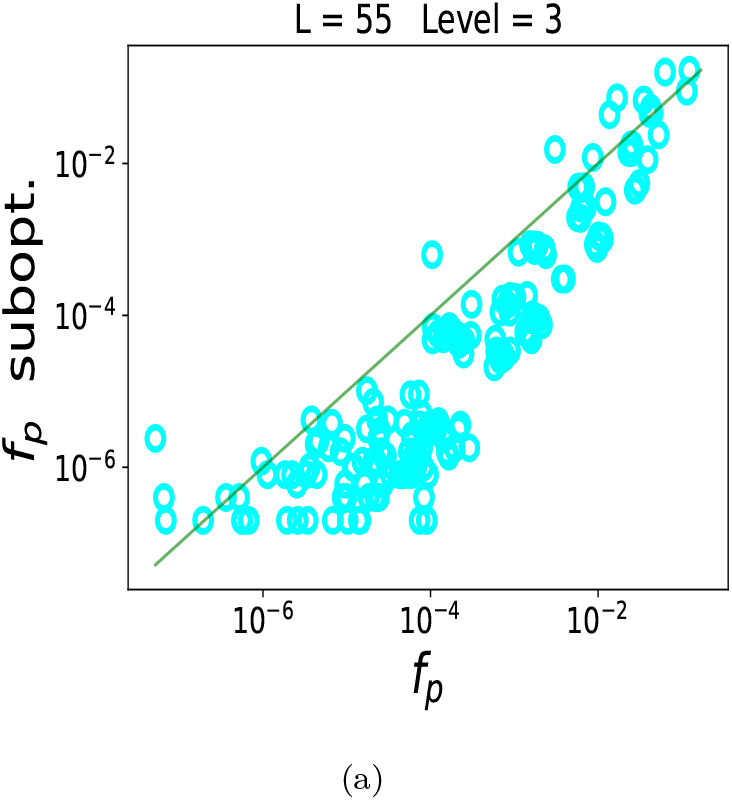
There is a high correlation between the frequencies of RNA shapes when using a single shape for each sequence (x-axis), and when incorporating suboptimal structures (y-axis).

Figure (S5) was obtained using the *L* = 55 level 3 data from the main text (5 × 10^6^ samples) to specify *f*_*p*_, and 1000 random sequences with their many respective suboptimal structures and shapes to specify *f*_*p*_ *subopt*.. All suboptimal structures up to 10kT or 6kcal/mol were incorporated for each sequence (note that the number of different suboptimal structures grows very large for large energy gaps, but this is compensated by small probabilities for larger energy gaps).

The correlation in Figure (S5) is high at 0.90 (p-val*<* 10^*−*6^) which indicates that whether we study the frequencies of MFE shapes as defined by *f*_*p*_ (as employed in the maintext) or defined by *f*_*p*_ *subopt*., we would get qualitatively similar results. Nevertheless there are small differences, and these may vary from structure to structure, suggesting that this direction of research is promising for future investigations.

## References

[1] J. M. Smith, R. Burian, S. Kauffman, P. Alberch, J. Campbell, B. Goodwin, R. Lande, D. Raup, and L. Wolpert, The Quarterly Review of Biology 60, 265 (1985).

[2] G. P. Wagner and L. Altenberg, Evolution 50, 967 (1996).

[3] S. J. Gould, The structure of evolutionary theory (Harvard University Press, 2002).

[4] A. Wagner, Arrival of the Fittest: Solving Evolution’s Greatest Puzzle (Penguin, 2014).

[5] K. Laland, G. A. Wray, and H. E. Hoekstra, Nature 514, 161 (2014).

[6] D. M. McCandlish and A. Stoltzfus, The Quarterly review of biology 89, 225 (2014).

[7] A. C. Love, Conceptual change in biology, Vol. 307 (Springer, 2015).

[8] D. Charlesworth, N. H. Barton, and B. Charlesworth, Proceedings of the Royal Society B: Biological Sciences 284, 20162864 (2017).

[9] A. Stoltzfus, arXiv preprint 1805.06067 (2018).

[10] T. Uller, A. P. Moczek, R. A. Watson, P. M. Brakefield, and K. N. Laland, Genetics 209, 949 (2018).

[11] E. I. Svensson and D. Berger, Trends in ecology & evolution 34, 422 (2019).

[12] T. Uller and K. Laland, Evolutionary causation: biological and philosophical reflections, Vol. 23 (the MIT press, 2019).

[13] D. Jablonski, Evolution & development 22, 103 (2020).

[14] A. A. Louis, Studies in History and Philosophy of Science Part C: Studies in History and Philosophy of Biological and Biomedical Sciences 58, 107 (2016).

[15] D. M. Raup, Journal of Paleontology 40, 1178 (1966).

[16] G. McGhee, The geometry of evolution: adaptive landscapes and theoretical morphospaces (Cambridge University Press, 2007).

[17] S. E. Ahnert, Journal of The Royal Society Interface 14, 20170275 (2017).

[18] S. Manrubia, J. A. Cuesta, J. Aguirre, S. E. Ahnert, L. Altenberg, A. V. Cano, P. Catalán, R. Diaz-Uriarte, S. F. Elena, J. A. García-Martín, et al., Physics of Life Reviews 38, 55 (2021).

[19] J. S. Mattick and I. V. Makunin, Human molecular genetics 15, R17 (2006).

[20] W. Gilbert, Nature 319, 618 (1986).

[21] E. P. Consortium et al., Nature 489, 57 (2012).

[22] A. F. Palazzo and E. S. Lee, Frontiers in genetics 6, 2 (2015).

[23] B. C. Thiel, C. Flamm, and I. L. Hofacker, Emerging Topics in Life Sciences 1, 275 (2017).

[24] Z. Miao, R. W. Adamiak, M. Antczak, M. J. Boniecki, J. M. Bujnicki, S.-J. Chen, C. Y. Cheng, Y. Cheng, F.-C. Chou, R. Das, et al., RNA, 075341 (2020).

[25] P. Schuster, W. Fontana, P. Stadler, and I. Hofacker, Proceedings: Biological Sciences 255, 279 (1994).

[26] R. Lorenz, S. H. Bernhart, C. H. Zu Siederdissen, H. Tafer, C. Flamm, P. F. Stadler, and I. L. Hofacker, Algorithms for molecular biology 6, 26 (2011).

[27] I. Hofacker, W. Fontana, P. Stadler, L. Bonhoeffer, M. Tacker, and P. Schuster, Monatshefte für Chemie/Chemical Monthly 125, 167 (1994).

[28] W. Fontana, BioEssays 24, 1164 (2002).

[29] A. Wagner, Robustness and evolvability in living systems (Princeton University Press Princeton, NJ:, 2005).

[30] R. Knight, H. De Sterck, R. Markel, S. Smit, A. Oshmyansky, and M. Yarus, Nucleic Acids Research 33, 5924 (2005).

[31] S. Smit, M. Yarus, and R. Knight, RNA 12, 1 (2006).

[32] M. Stich, C. Briones, and S. C. Manrubia, Journal of Theoretical Biology 252, 750 (2008).

[33] T. Jorg, O. Martin, and A. Wagner, BMC bioinformatics 9, 464 (2008).

[34] M. Cowperthwaite, E. Economo, W. Harcombe, E. Miller, and L. Meyers, PLoS computational biology 4, e1000110 (2008).

[35] J. Aguirre, J. M. Buldú, M. Stich, and S. C. Manrubia, PloS one 6, e26324 (2011).

[36] S. Schaper and A. A. Louis, PloS one 9, e86635 (2014).

[37] A. Wagner, The Origins of Evolutionary Innovations: A Theory of Transformative Change in Living Systems (Oxford University Press, 2011).

[38] K. Dingle, S. Schaper, and A. A. Louis, Interface focus 5, 20150053 (2015).

[39] S. F. Greenbury, S. Schaper, S. E. Ahnert, and A. A. Louis, PLoS computational biology 12, e1004773 (2016).

[40] C. G. Oliver, V. Reinharz, and J. Waldispühl, RNA 25, 1579 (2019).

[41] J. A. García-Martín, P. Catalán, S. Manrubia, and J. A. Cuesta, EPL (Europhysics Letters) 123, 28001 (2018).

[42] M. Weiß and S. E. Ahnert, Journal of The Royal Society Interface 15, 20170618 (2018).

[43] T. Mituyama, K. Yamada, E. Hattori, H. Okida, Y. Ono,G. Terai, A. Yoshizawa, T. Komori, and K. Asai, Nucleic Acids Research 37, D89 (2009).

[44] W. Fontana, D. A. Konings, P. F. Stadler, and P. Schuster, Biopolymers 33, 1389 (1993).

[45] R. Giegerich, B. Voß, and M. Rehmsmeier, Nucleic Acids Research 32, 4843 (2004).

[46] Nucleic acids research 49, D212 (2021).

[47] D. H. Mathews, M. D. Disney, J. L. Childs, S. J. Schroeder, M. Zuker, and D. H. Turner, Proceedings of the National Academy of Sciences 101, 7287 (2004).

[48] J. S. Reuter and D. H. Mathews, BMC bioinformatics 11, 1 (2010).

[49] N. R. Markham and M. Zuker, in Bioinformatics (Springer, 2008) pp. 3–31.

[50] S. Janssen and R. Giegerich, Bioinformatics 31, 423 (2015).

[51] M. E. Nebel and A. Scheid, Theory in Biosciences 128, 211 (2009).

[52] W. A. Lorenz, Y. Ponty, and P. Clote, Journal of Computational Biology 15, 31 (2008).

[53] I. Kalvari, J. Argasinska, N. Quinones-Olvera, E. P. Nawrocki, E. Rivas, S. R. Eddy, A. Bateman, R. D. Finn, and A. I. Petrov, Nucleic acids research 46, D335 (2018).

[54] I. Kalvari, E. P. Nawrocki, J. Argasinska, N. Quinones-Olvera, R. D. Finn, A. Bateman, and A. I. Petrov, Current protocols in bioinformatics 62, e51 (2018).

[55] F. B. Salisbury, Nature 224, 342 (1969).

[56] J. M. Smith, Nature (1970).

[57] F. Jacob, Science 196, 1161 (1977).

[58] R. Neme, C. Amador, B. Yildirim, E. McConnell, and D. Tautz, Nature Ecology & Evolution 1, 1 (2017).

[59] D. J. Begun, H. A. Lindfors, A. D. Kern, and C. D. Jones, Genetics 176, 1131 (2007).

[60] D. Tautz and T. Domazet-Lošo, Nature Reviews Genetics12, 692 (2011).

[61] B. A. Wilson, S. G. Foy, R. Neme, and J. Masel, Nature Ecology & Evolution 1, 1 (2017).

[62] M. de la Peña and I. García-Robles, RNA 16, 1943 (2010).

[63] L. Yampolsky and A. Stoltzfus, Evolution & Development 3, 73 (2001).

[64] A. Stoltzfus and D. M. McCandlish, Molecular biology and evolution 34, 2163 (2017).

[65] A. V. Cano and J. L. Payne, PLoS computational biology 16, e1008296 (2020).

[66] M. Lynch, Proceedings of the National Academy of Sciences 104, 8597 (2007).

[67] P. Catalán, S. Manrubia, and J. A. Cuesta, Journal of the Royal Society Interface 17, 20190843 (2020).

[68] C. Wilke, J. Wang, C. Ofria, R. Lenski, and C. Adami, Nature 412, 331 (2001).

[69] G. Valle-Pérez, C. Q. Camargo, and A. A. Louis, arXiv preprint 1805.08522 (2018).

[70] C. Mingard, J. Skalse, G. Valle-Pérez, D. Martínez-Rubio, V. Mikulik, and A. A. Louis, arXiv preprint 1909.11522 (2019).

[71] L. Bottou, F. E. Curtis, and J. Nocedal, Siam Review 60, 223 (2018).

[72] C. Mingard, G. Valle-Pérez, J. Skalse, and A. A. Louis, Journal of Machine Learning Research 22, 1 (2021).

[73] K. Gomez, J. Bertram, and J. Masel, Proceedings of the Royal Society B 287, 20201503 (2020).

[74] C. Tuerk and L. Gold, Science 249, 505 (1990).

[75] A. D. Ellington and J. W. Szostak, nature 346, 818 (1990).

[76] M. M. Vu, N. E. Jameson, S. J. Masuda, D. Lin, R. Larralde-Ridaura, and A. Lupták, Chemistry & biology 19, 1247 (2012).

[77] K. Salehi-Ashtiani and J. Szostak, Nature 414, 82 (2001).

[78] E. Mayr, Science (New York, NY) 134, 1501 (1961).

[79] K. N. Laland, K. Sterelny, J. Odling-Smee, W. Hoppitt, and T. Uller, Science 334, 1512 (2011).

[80] R. Scholl and M. Pigliucci, Biology & Philosophy 30, 653 (2015).

[81] W. Arthur, Evolution & development 3, 271 (2001).

[82] G. B. West and J. H. Brown, Journal of experimental biology 208, 1575 (2005).

[83] K. Dingle, C. Q. Camargo, and A. A. Louis, Nature communications 9, 761 (2018).

[84] D. Thompson, On Growth and Form (Cambridge University Press, 1942).

[85] T. Schlick and A. M. Pyle, Biophysical journal 113, 225 (2017).

[86] E. Rivas, J. Clements, and S. R. Eddy, Nature methods 14, 45 (2017).

[87] E. Rivas, J. Clements, and S. R. Eddy, Bioinformatics 36, 3072 (2020).

[88] I. G. Johnston, K. Dingle, S. F. Greenbury, C. Q. Camargo, J. P. K. Doye, S. E. Ahnert, and A. A. Louis, bioRxiv (2021), 10.1101/2021.07.28.454038.

[89] S. Ahnert, I. Johnston, T. Fink, J. Doye, and A. Louis, Physical Review E 82, 026117 (2010).

[90] I. G. Johnston, S. E. Ahnert, J. P. Doye, and A. A. Louis, Physical Review E 83, 066105 (2011).

[91] S. F. Greenbury, I. G. Johnston, A. A. Louis, and S. E. Ahnert, Journal of The Royal Society Interface 11, 20140249 (2014).

## References

[1] R. Giegerich, B. Voß, and M. Rehmsmeier, Nucleic Acids Research 32, 4843 (2004).

[2] I .Kalvari, J. Argasinska, N. Quinones-Olvera, E. P. Nawrocki, E. Rivas, S. R. Eddy, A. Bateman, R. D. Finn, and A. I. Petrov, Nucleic acids research 46, D335 (2018).

[3] I. Kalvari, E. P. Nawrocki, J. Argasinska, N. Quinones-Olvera, R. D. Finn, A. Bateman, and A. I. Petrov, Current protocols in bioinformatics 62, e51 (2018).

[4] J. Waldispühl and Y. Ponty, in Research in Computational Molecular Biology, edited by V. Bafna and S. C. Sahinalp (Springer Berlin Heidelberg, Berlin, Heidelberg, 2011) pp. 501–515.

[5] K. Dingle, S. Schaper, and A. A. Louis, Interface focus 5, 20150053 (2015).

